# A multi-scale segmentation-free self-supervised AI model to characterize the heterogeneity of the brain tumor microenvironment

**DOI:** 10.1101/2025.02.20.639309

**Authors:** Sam Sterling, Jimin Tan, Hortense Le, Danielle Share, Yi Ban, Matija Snuderl, Aristotelis Tsirigos

## Abstract

Brain tumors affect about 1 million people in the U.S., with aggressive types like glioblastoma having very low survival rates due to complex tumor biology and the protective blood-brain barrier. Current treatments are limited in effectiveness, and our understanding of brain tumor biology remains incomplete. High dimensional multiplexed imaging has enabled us to better understand the tumor microenvironment (TME); however, analyses typically rely on cell segmentation, which is error-prone, may discard useful context outside the cell boundary, and neglects complex tissue-wide features. To address this limitation, we developed a segmentation-free, self-supervised representation learning framework that enables us to train directly on multiplexed images using masked image modeling. We used this approach to analyze 389 imaging mass cytometry images from 185 brain tumor patients. To study tissue-wide features, we first trained our model on 64×64 micron tiles capturing neighborhoods of 10-20 cells, which we termed local tumor microenvironments (LTMEs). To further characterize these LTMEs, we trained our model on 16×16 micron tiles centered on individual cells in our dataset, so that each tile captures a single cell and its surrounding area, which we termed single-cell microenvironments (SCMEs). This multi-scale, self-supervised approach enables a detailed analysis of the heterogeneity within the brain TME, examining single cells in their spatial context. In addition to validating known findings, we identified a novel LTME in GBM patients, composed primarily of tumor cells and a few B and T cells, which strongly correlated with increased survival. By analyzing these B cells with our SCME model, we found they were distinct from other GBM B cells, and higher concentrations of these B cells were linked to improved survival. In conclusion, our study introduces a multi-scale, segmentation-free, self-supervised machine learning model that provides unprecedented insights into brain TMEs, enabling discovery of previously unrecognized cell interactions and spatial features that are predictive of patient survival.

## Introduction

In the United States, up to 1 million individuals are living with a brain tumor. An estimated 90,000 new cases of primary brain tumor are diagnosed each year, the most common of which are astrocytomas, a glioma that originates from glial cells called astrocytes. Glioblastoma (GBM) is the most common malignant brain tumor and has a median survival of just one year. Additionally, between 21,000 to 400,000 cases of brain metastases are diagnosed per year, which have a survival rate of just 6 to 10 months.^1–5^ Brain tumors are characterized by highly heterogeneous, spatially complex microenvironments, defined by factors such as immune privilege and specialized cell interactions. The blood brain barrier, intended to facilitate the brain’s stable internal environment, greatly reduces the effectiveness of many cancer treatments including chemotherapy.^6^ Immune activity is greatly restricted, as the blood-brain barrier limits the movement of immune cells such as T cells and B cells into the brain, and the brain’s immune-privileged status dampens immune activity to protect brain tissue from inflammation.^7^ Tumor cells can also reprogram immune cells to secrete factors that promote tumor growth and survival.^8^ Current neurosurgical or radiotherapeutic treatment options significantly impair quality of life without substantially prolonging survival, and clinical trials have yet to yield an effective therapy for most brain tumors, partially due to our incomplete understanding of brain tumor biology.^3,9^ Brain tumor research also receives limited funding and research due to its relative rarity among cancers, and patients are often excluded from phase 1 clinical trials due to concerns around potential complications.^9,10^

Recent advances in multiplexed imaging techniques, such as imaging mass cytometry (IMC), have made it possible to capture up to 40 protein markers in spatial detail, offering new insights into how immune cells infiltrate tumors and how vascular changes contribute to tumor progression.^11^ Traditional tools for analyzing these complex datasets, such as histoCAT, rely on cell segmentation.^12^ Because segmentation-based approaches focus on individual cells, they may fail to capture the full complexity of spatial interactions in heterogeneous brain tumors and limit our understanding of tissue-wide features that drive tumor progression.

A recent study conducted by the Quail-Walsh Lab at McGill University used IMC to map the immune landscapes of primary and metastatic brain tumors and defined spatial cellular neighborhoods formed by clustering frequency vectors of the nearest neighboring cells around each cell. This study identified a novel population of MPO+ macrophages associated with long term survival.^6^ Using the imaging data from this study, we applied a segmentation-free, self-supervised model to characterize known and novel features of the brain tumor microenvironment. We adopted a multi-scale approach, analyzing 64×64 micron tiles that capture neighborhoods of 10-20 cells, which we term local tumor microenvironments (LTMEs), as well as 16×16 micron tiles centered on individual cells and their surrounding area, which we term single-cell microenvironments (SCMEs). We identified 40 LTME types by clustering and found that one (L2), enriched with tumor cells and some B and T cells, predicts increased survival in GBM patients. Moreover our SCME analysis reveals B cells present in the L2 LTME express distinct markers from other B cells in GBM and are associated with survival differences.

## Results

### A multi-scale self-supervised AI model to study the TME heterogeneity of brain tumors

The dataset consists of 389 IMC images from 139 patients with high-grade glioma, and 46 patients with metastatic brain cancer. Of the 139 patients with high-grade gliomas, 118 were classified as IDH wt (glioblastoma), 14 as IDH mutant (astrocytoma), and 7 as unknown IDH status. The stage, MGMT status, and post-surgical treatment status are presented in Table 1 below. The data also included samples from 46 brain metastasis patients, consisting of patient-matched images from both the core of the metastatic lesion and the surrounding brain tissue interface, which are presented in Table 2.

**Table 1:** Glioma patients by IDH status, stage, MGMT promoter status, and post-surgical treatment.

| IDH status | Stage |  | MGMT promoter status |  |  | Post-surgical treatment |  | Total |
| --- | --- | --- | --- | --- | --- | --- | --- | --- |
|  | Primary | Recurrent | um | m | unknown | SOC | non-SOC |  |
| Wild-type (glioblastoma) | 106 | 12 | 54 | 62 | 2 | 99 | 19 | 118 |
| Mutant (astrocytoma) | 10 | 4 | 6 | 8 | 0 | 13 | 1 | 14 |
| Unknown | 7 | 0 | 1 | 4 | 1 | 7 | 0 | 7 |

**Table 2:** Brain metastases patients by primary tumor site and radiation treatment.

| Primary tumor site | Treated with radiation | Not treated with radiation | Unknown treatment | Total |
| --- | --- | --- | --- | --- |
| Breast | 9 | 2 | 0 | 11 |
| Lung | 18 | 2 | 1 | 20 |
| Melanoma | 5 | 1 | 2 | 8 |
| Ovary | 2 | 0 | 0 | 2 |
| Colorectal | 1 | 0 | 0 | 1 |
| Gastro-intestinal | 2 | 0 | 0 | 2 |
| Bladder | 1 | 1 | 0 | 2 |

To generate LTMEs we first performed masking on the original images to identify areas with high pixel intensity and then used these regions to generate 25,671 64-micron square tiles each containing around 10-20 cells. Next, we generated 16-micron square tiles centered on all 923,960 cells in our dataset, so that each tile captures a single cell and its surrounding area, which we termed single-cell microenvironments (SCMEs). We experimented with masking the area outside the tile boundary but ultimately decided to keep it as it improved model performance, likely because our model was able to learn from the additional context.

We used our self-supervised representation framework to train two separate models, one for LTME tiles and another for SCME tiles. Our model consists of a vision transformer encoder and decoder, which processes each tile as a series of 5 micron square patches. The encoder processes 25% of the patches, and the decoder is asked to reconstruct the remaining 75% of the masked patches. The unmasked images are then fed to the trained model in order to create feature embeddings, which we visualize with a UMAP (**Fig. 1**). For the LTME model we used all 25,671 LTME tiles. However, to limit computational time we used a representative sample of 230,953 SCME tiles for training the SCME model, by selecting up to 100 SCMEs of each cell type from each image.

**Fig. 1:**
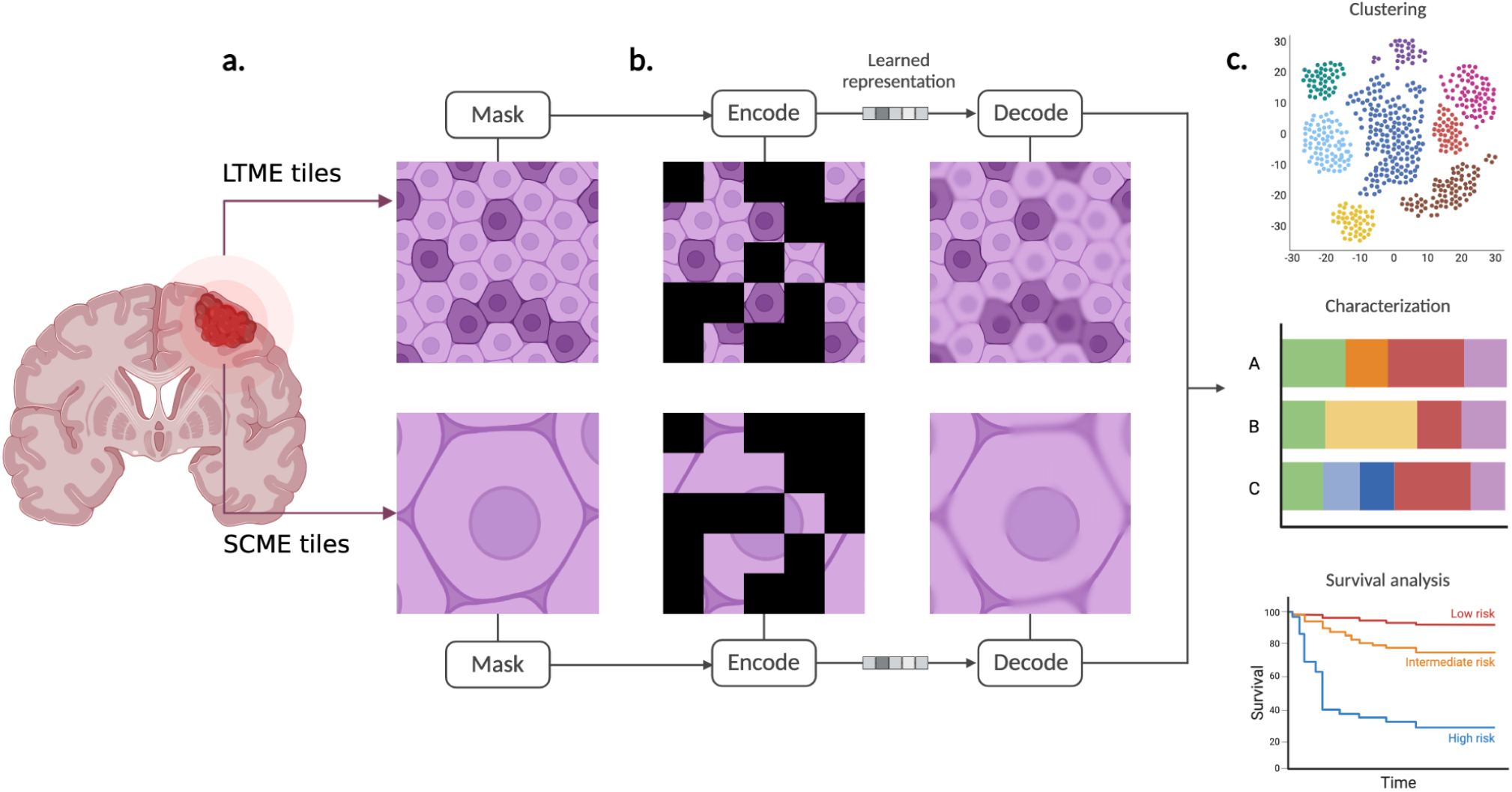
Multi-scale self-supervised learning framework. **a)** Tumor slides are divided into 64-micron square LTME tiles representing local neighborhoods of 10-20 cells, as well as 16-micron square SCME tiles centered on individual single cells. **b)** We apply masked image modeling to train our model using self-supervised masked image learning, in which 75% of tiles are randomly masked and used as reconstruction targets for the decoder. **c)** A feature embedding is generated for each tile and we cluster feature embeddings to perform downstream analysis, including association of cluster composition with clinical outcomes.

For both models, we used channel-wise normalization across our markers to ensure consistent scaling. Image augmentation was applied to mimic real-world biological variations in the data; this comprised random rotations (0 to 180 degrees), horizontal flips, random cropping (scaling from 50 to 100%), and channel intensity augmentation. For SCME tiles, our random cropping scaled from 80 to 100% to ensure that in most cases the entire cell remains inside the tile.

We applied k-means clustering to identify 40 LTME and 40 SCME clusters. For each patient, we calculate their tile concentration (TC) for each LTME, which we define as the number of tiles a patient has for a specific LTME cluster divided by their total number of tiles. For each LTME cluster, we are then able to calculate median TC across all patients. Patients were then divided into high and low TC groups based on whether their TC for an LTME or SCME was above or below the median. Survival analysis was performed to evaluate the association between high and low TC groups and patient survival for each LTME cluster.

### Self-supervised representation learning identifies a diverse landscape of LTMEs in brain tumors

The model was first trained on all LTME tiles with masking, with the original images used as reconstruction targets. After training, the unmasked images are fed into the encoder to generate embeddings for all LTME tiles, which we visualized with a UMAP (**Fig. 2a**). As expected, LTME tiles from glioma and BrM patients form two distinct clusters, and BrM LTMEs cluster based on the primary tumor site (**Fig. 2b**). We used k-means to group our LTME feature embeddings into clusters, and tested using 10, 20, 30, 40, 50, and 60 centroids for clustering. We ultimately settled on 40 centroids, as the cell and marker composition chart for each cluster showed that 40 clusters yielded a diverse set of clusters with minimal redundant clusters. By plotting cell composition and marker expression for each LTME (**Fig. 3**), we observed that most of our LTMEs consisted of tumor cells with a few immune cells, reflecting the heterogeneous and immune-deficient nature of brain tumors. We then utilized the LTME clusters to characterize the heterogeneity of the patients’ tumors. To this end, we clustered patients into 9 groups based on their LTME composition, and created a heatmap to visualize the relationship between patient clusters, clinical variables, and LTME enrichment (**Fig. 4**). As expected, BrM and glioma patients cluster into separate groups and are enriched for different LTMEs. These results highlight the diversity and complexity of the LTME landscape both within a specific tumor and across patients. In the following sections, we will investigate in detail several of these LTMEs in terms of their cell type composition and their links to clinical outcomes.

**Fig. 2:**
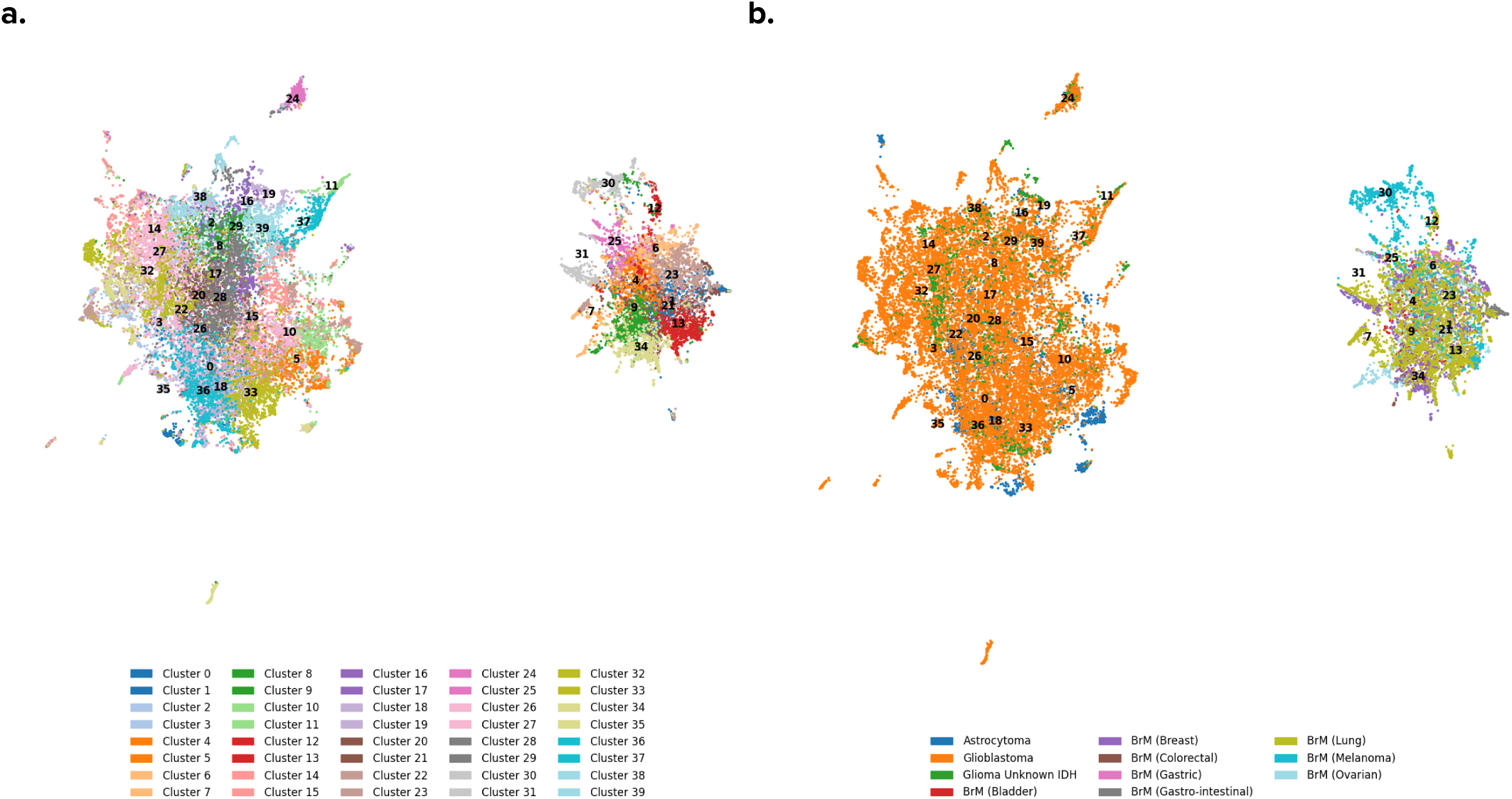
Self-supervised representation learning of local cell neighborhoods identifies clusters of LTMEs. **a.** UMAP of LTME feature embeddings with 40 centroids, where each dot represents an LTME tile, and numbers represent the center of that cluster. **b.** UMAP of LTME embeddings colored by tumor type: as expected, we see a clearer separation between our glioma and BrM tiles compared to our SCME tiles, as glioma and BrM tumors have distinct microenvironments.

**Fig. 3:**
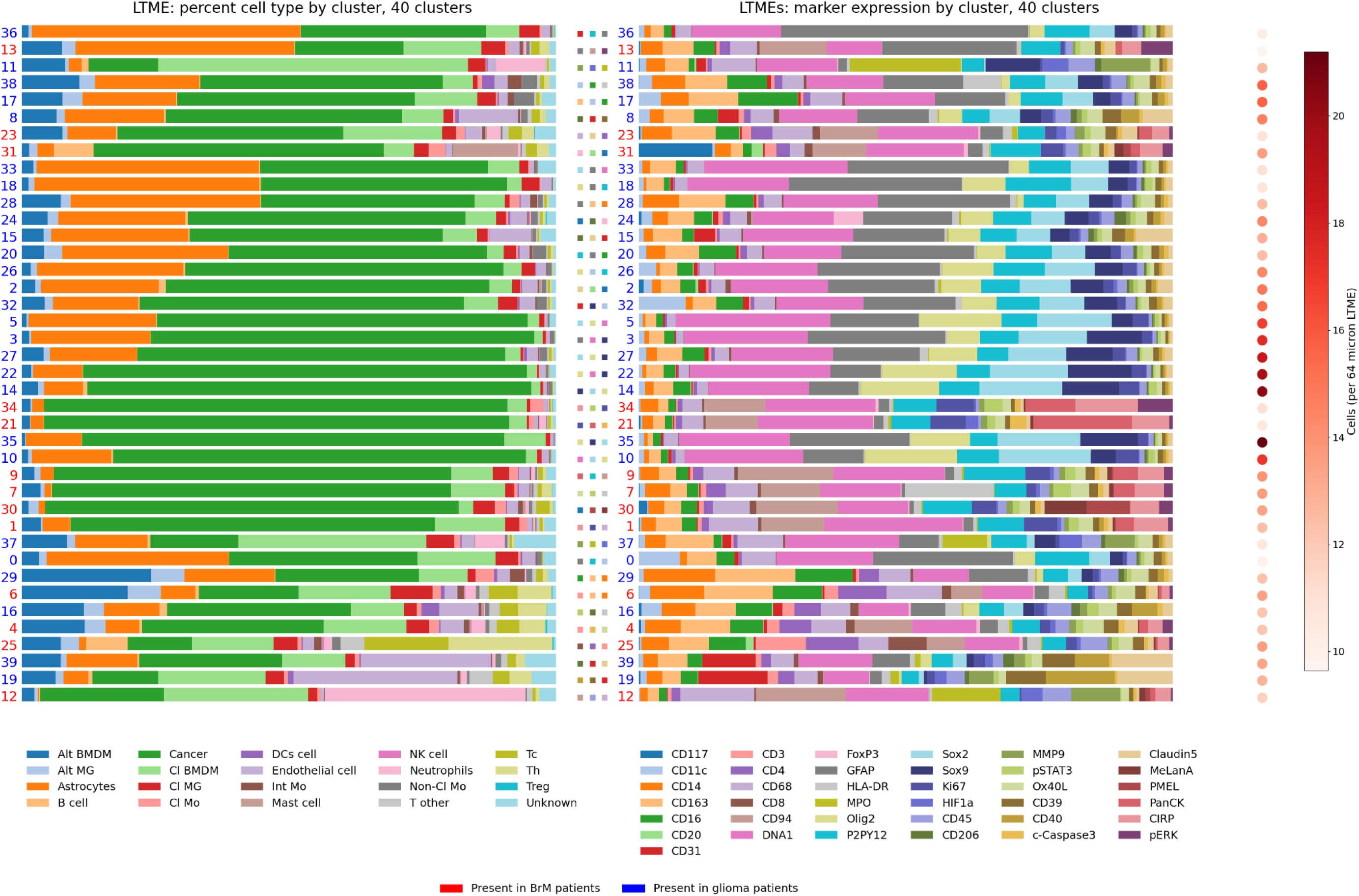
Bar charts show the normalized cell composition (left) marker composition (right) and density for each LTME cluster. Blue and **red** cluster numbers indicate at least 90% of the LTMEs in this cluster are present in glioma or BrM samples respectively.

**Fig. 4:**
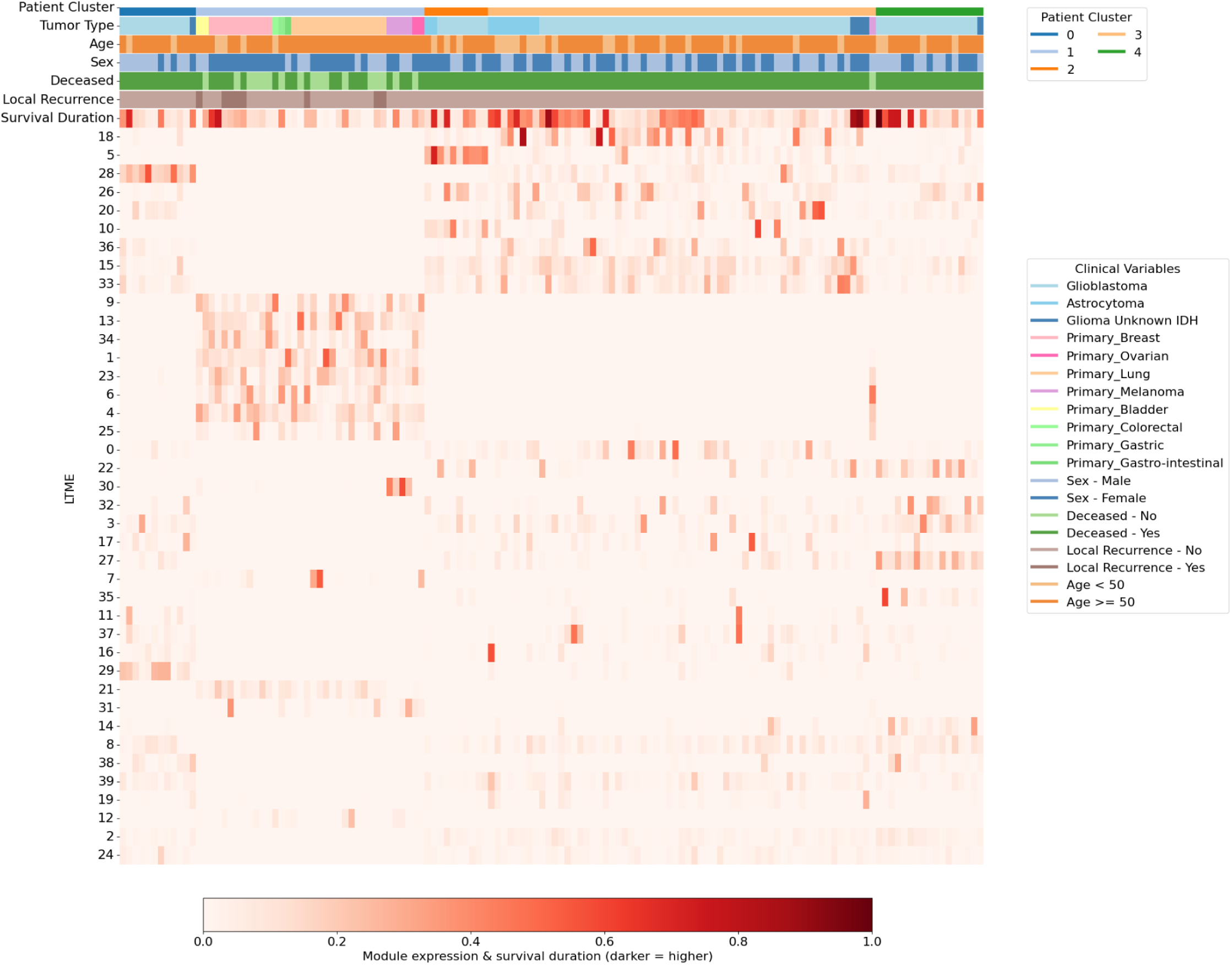
Heatmap showing correlations between patient clusters, clinical variables, and LTME cluster composition.

### Self-supervised representation learning on single-cell microenvironments (SCMEs)

In parallel, we applied the trained SCME model to the SCME tiles, obtained embeddings for each tile, generated 40 clusters using k-means clustering, and visualized the result by a UMAP. To validate our SCME clusters, we visualized marker expression by cluster, and compared it to marker expression for the annotated cell types in the published dataset (**Suppl. Fig. 2**). Next, we plotted each feature embedding on a UMAP colored by cell- and tumor-type, and noted that cells cluster by both cell type and tumor type (primary versus metastases) (**Suppl. Fig. 3**). Brain metastases cells also cluster by primary tumor, supporting research showing that the location of the primary tumor affects protein expressions in the metastatic tumor.^13^

### LTME cluster L2 is associated with increased survival in glioblastoma patients

To examine the relationship between LTMEs and clinical outcome, we split patients into high and low groups at the 50th, 25th, and 10th percentile of TC, plotted survival curves using Kaplan-Meier, calculated p-values with a log-rank test, and performed multiple comparisons corrections using the Benjamini-Hochberg false discovery rate method. To validate significant clusters further, we calculated the concordance index (c-index) to assess how well the model predicts the order of patient survival times, and performed multivariate Cox analysis to determine hazard ratios for each cluster.

Only one LTME had a significant association with survival: L2, predominantly observed in glioblastoma (IDH wt) patients, significantly predicts increased survival in our Kaplan-Meier analysis **(Fig. 5a)** and multivariate Cox Hazards analysis (coef = −15.07, p < 0.005). When splitting at the 50th percentile, patients in the high TC group survived an average of 1,175 days compared to 608 days for the low TC group. L2 also predicts increased survival in multivariate Cox hazards (p < 0.005). Our concordance index model, which uses L2 TC to rank patient survival, generated a c-index score of 0.623. While a c-index score of 0.623 typically does not indicate very high accuracy, predicting survival in GBM is notoriously difficult due to its aggressive nature and genetic variability. In a recent study, researchers used several independent prognostic factors to develop a nomogram in order to predict GBM survival outcomes, and achieved a c-index of 0.729.^14^ Controlling for MGMT methylation status, a predictive biomarker of survival in GBM patients^15^, did not affect L2’s ability to predict survival. We did not control for molecular subtypes or DNA methylation class, as that information was not available for our dataset.

**Fig. 5:**
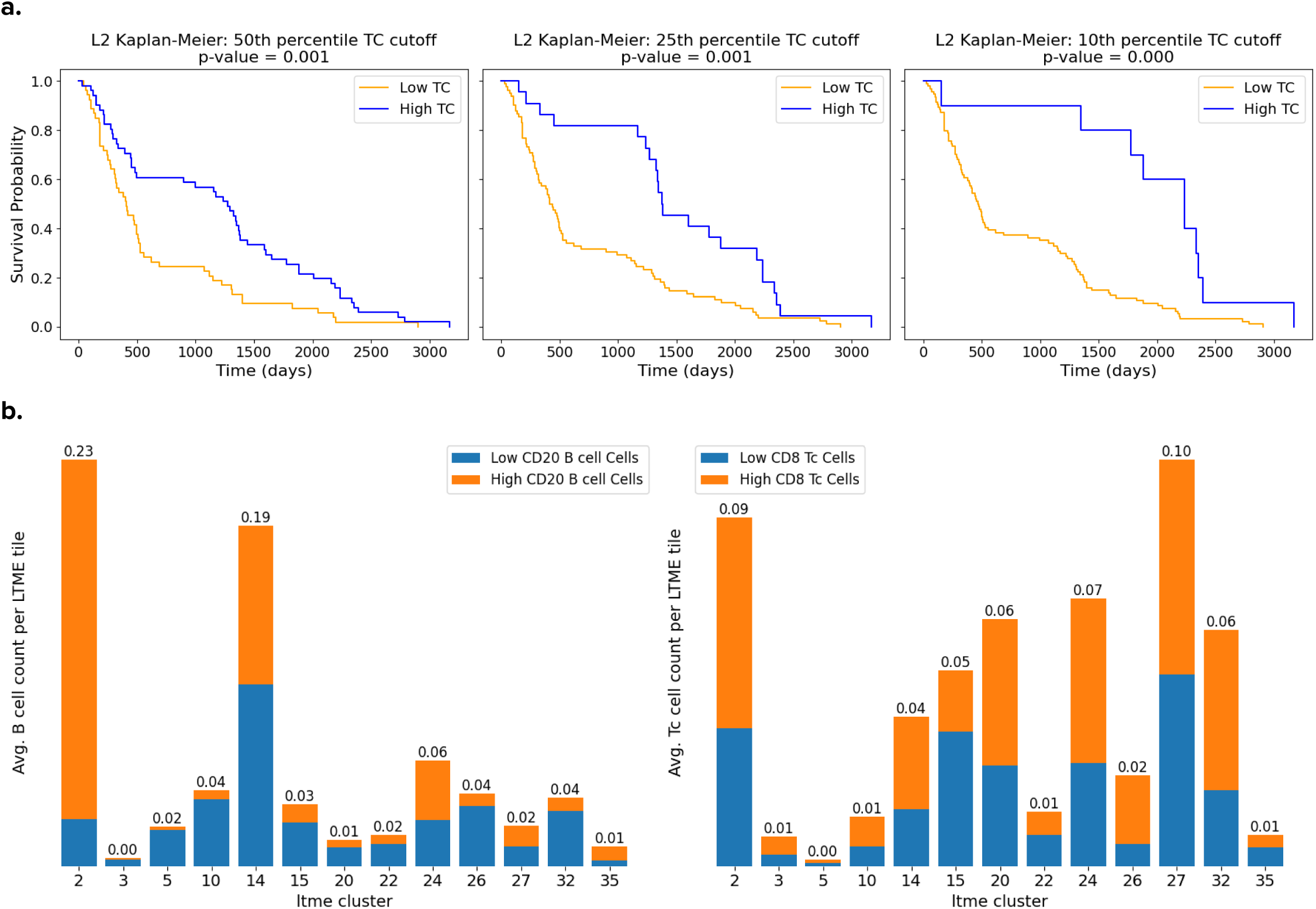
L2 LTME tiles contain high levels of CD20+ B cells and CD8+ T cells and predict increased survival. a) Kaplan Meier curves showing that L2 predicts increased survival given cutoffs at the 50th, 25th, and 10th percentile of TC. b) Compared with other glioma tumor LTME clusters, L2 is characterized by a higher average quantity of CD20+ B cells and CD8+ T cells.

In terms of cell composition, L2 closely resembles our other glioma tumor LTME clusters (defined as clusters with at least 90% of tiles from glioma patients and over 50% tumor cells), but unlike most glioma tumor LTME clusters, it contains a small quantity of B cells, which are a rare feature in GBM. We also observe that relative to other glioma LTMEs consisting primarily of tumor cells, L2 consists of B cells with higher CD20 expression, and T cells with higher CD8 expression **(Fig. 5b)**. While L2 does show interactions between B cells, T cells, dendritic cells, and endothelial cells, we did not observe any evidence of structured tertiary lymphoid structures, which are organized immune aggregates with a structural and functional resemblance to secondary lymphoid organs that can sometimes lead to improved patient outcomes **(Fig. 6)**.^16^

**Fig. 6:**
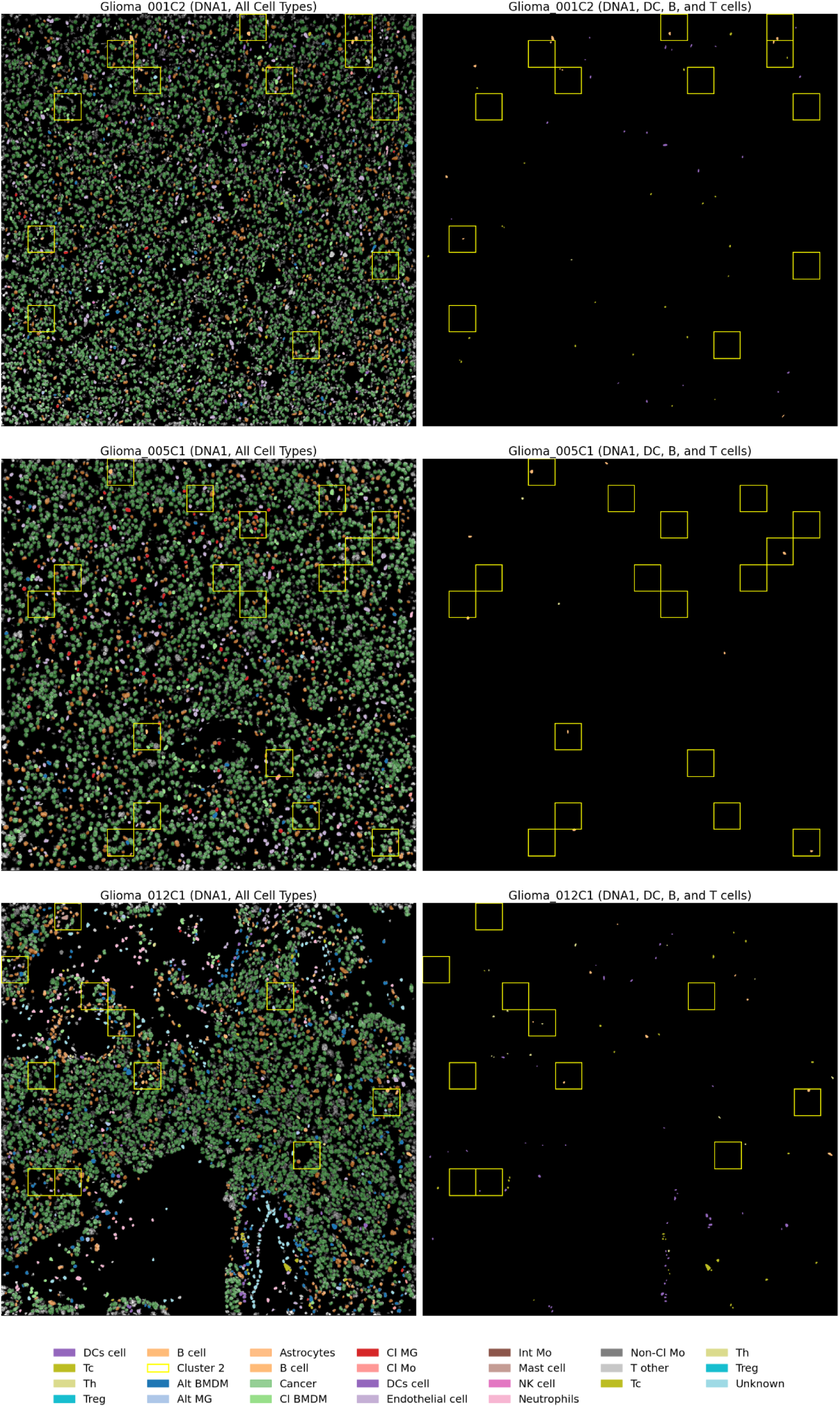
Visual analysis of L2 reveals the presence of immune cells but not structured tertiary lymphoid structures.

### SCME analysis reveals a context-specific role of B cells associated with contrasting patient outcomes

In many cancers, a high density of tumor infiltrating B cells is associated with increased survival, in part due to their role as antigen-presenting cells.^17^ But in glioblastoma, B cells are typically suppressed from maturing into active immune cells due to inhibitory checkpoint molecules and TGFβ signaling from tumor and myeloid cells. These suppressed B cells, which can include regulatory B cells, or Bregs, can predict aggressive tumor growth and poor prognosis. (16)

To ask if L2 contains a novel subtype of B cells, we clustered B cell SCME tiles from glioma patients into two clusters as suggested by our silhouette plot **(Suppl. Fig. 4)**. Using a web app that allows us to understand the spatial orientation and marker expression of LTMEs and cells **(Suppl. Fig. 7)**, we observed that Cluster 0 B cells are commonly found in perivascular regions and are characterized by higher expression of CD20 and CD45 **(Fig. 7b)**. Cluster 1 B cells, which represent the majority of B cells identified in our dataset, are characterized by low CD20 expression and are located further away from perivascular regions **(Suppl. Fig. 6).**

**Fig. 7:**
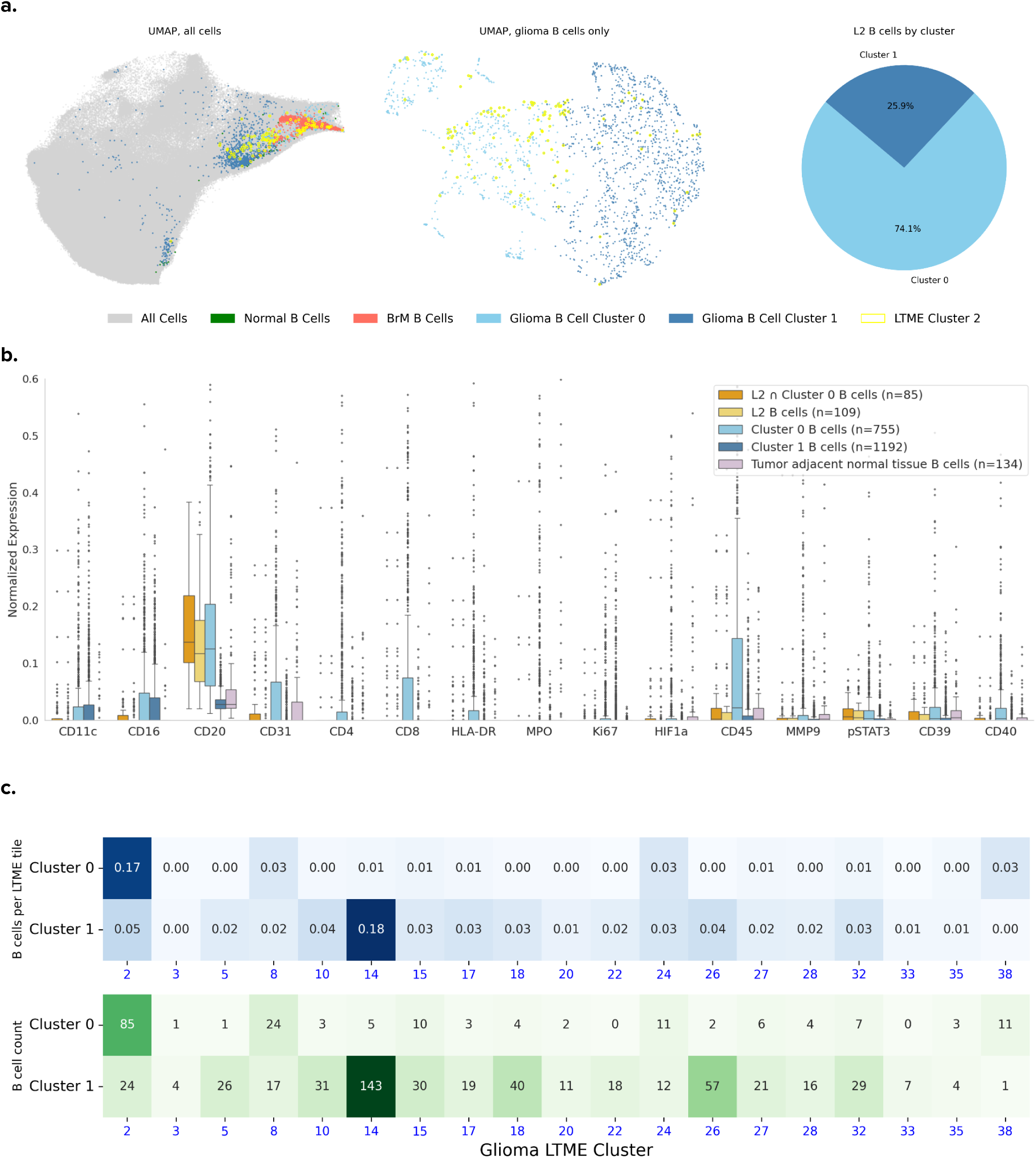
B cell SCMEs separate into 2 clusters that predict different survival times. **a)** UMAPs showing how glioma B cell SCME clusters differ from each other and BrM cells. Pie chart shows that most L2 B cells (circled in yellow) are in glioma B cell cluster 0. **b)** Box plots comparing marker expression for L2 B cells against Cluster 0, 1, and 2 B cells. L2 and C0 B cells are both characterized by higher CD20 expression. C0 B cells appear to express CD31, but this is likely “noise” caused by the proximity of infiltrating C0 B cells to endothelial cells, as SCME tiles can overlap and sometimes contain multiple cells. **c)** Heatmap showing B cell percentages and counts between the two clusters for each glioma LTME, showing overlap between L2 and C0 B cells.

Survival analysis reveals that the presence of Cluster 1 B cells predicts shortened survival in patients, while the presence of Cluster 0 B cells predicts long-term survival. **(Fig. 8a-b)** As expected, L2 primarily consists of Cluster 0 B cells **(Fig. 7c)**. The association between Cluster 0 B cells and survival is no longer significant when excluding cells present in L2, and as L2 is only found in the tumor core away from perivascular regions, this may suggest that Cluster 0 B cells must migrate from the perivascular region and into the tumor in order to promote improved survival. The presence of CD8+ T cells in L2 **(Fig. 5b)** might indicate that Cluster 0 B cells induce T cell activation through their role as antigen-presenting cells, but we cannot confirm this as we do not observe meaningful levels of HLA-DR or CD40 in L2, and our data does not include other markers that would reflect T cell activation by B cells. Conversely, the presence of CD8+ T cells in L2 may mean that this is simply a region that is conducive to immune infiltration and T cell activation.

**Fig. 8:**
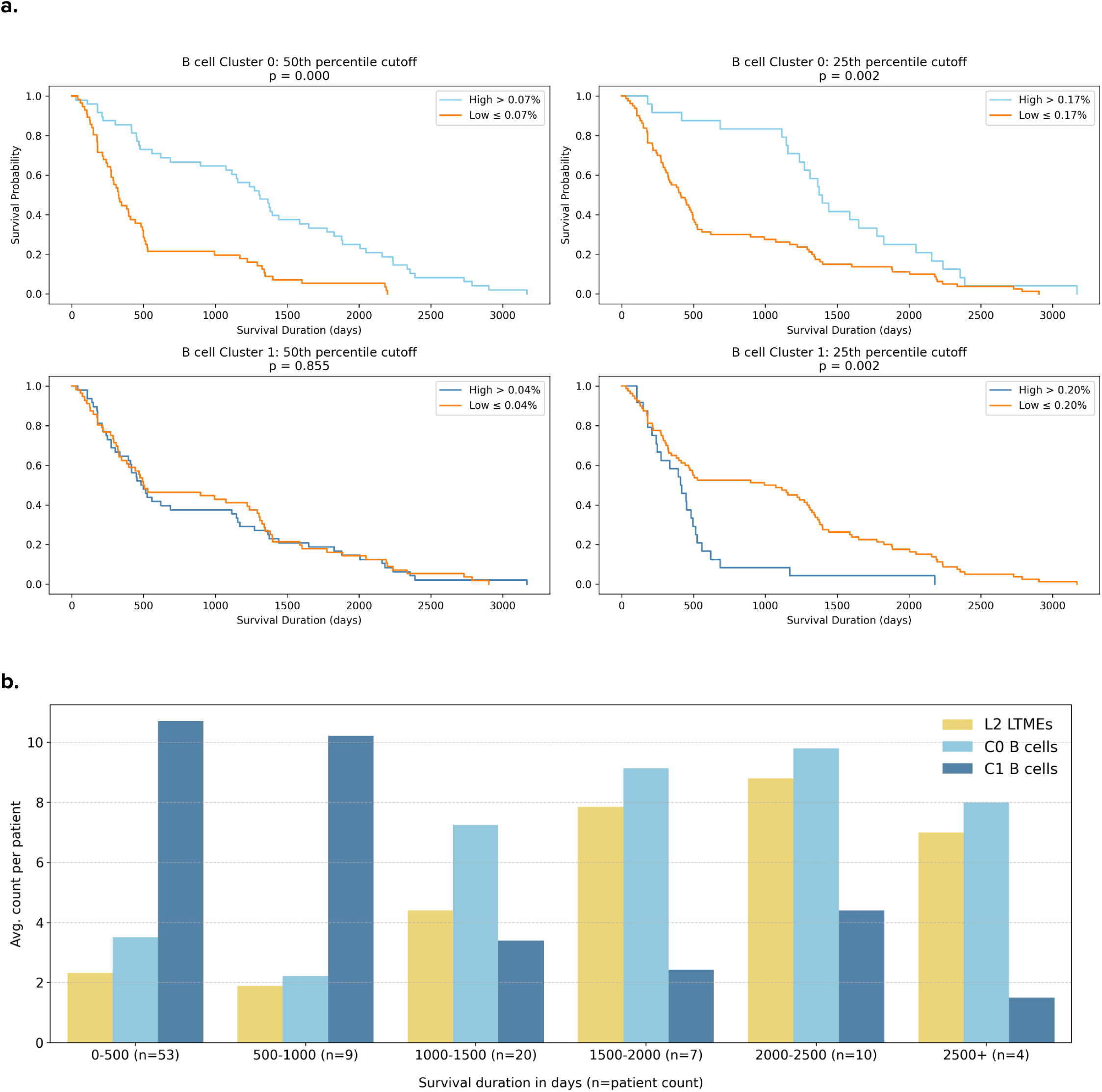
**a)** Kaplan-Meier curves showing that Cluster 0 strongly predicts improved survival at both the 50th and 25th percentile of TC, while Clusters 1 and 2 generally predict poorer survival. **b)** Bar chart showing average L2 LTME, as well as C0 and C1 B cell counts for all glioma patients, grouped by patient survival. C1 B cell prevalence tends to decrease with increased survival, while L2 and C0 B cell prevalence tend to increase.

Lastly, we analyzed three publicly available scRNA-seq and snRNA-seq datasets from glioblastoma patients. These datasets contained a total of 108,357 cells. Gene expression values were scaled from 0 to 1, and 398 cells were identified as B cells based on their expression of CD19, CD20, or CD79. B cells were divided into CD20^high^ and CD20^low^ groups using an expression threshold of 0.05, yielding a total of 38 CD20^high^ B cells. However, marker expression analysis of these two B cell types only served to confirm that a minority of B cells in GBM patients express CD20 at a much higher rate; CD20^high^ and CD20^low^ B cells were not significantly differentiated by expression of other marker types. **(Suppl. Fig. 5)**

## Discussion

In this paper, we leveraged a segmentation-free, self-supervised learning approach to identify novel spatial features of brain tumors, enabling us to identify cell subpopulations and tissue-wide features that predict survival. In doing so, we discovered a LTME (L2) that strongly correlates with long term survival in GBM IDH WT patients. By training our model on cells at the single-cell microenvironment (SCME) level, we identified two types of B cells primarily differentiated by their CD20 expression. Cluster 0 B cells have higher CD20 expression and predict increased survival, while Cluster 1 B cells have lower CD20 expression and predict decreased survival. As expected, L2 primarily consists of Cluster 0 B cells. While we are not able to confirm the functional or maturation state of these B cell types without additional markers, the contrasting correlation of these two B cell subtypes on survival represents a novel finding. Further characterization of these B cell subtypes and validation could enable their use as a predictive biomarker of survival in GBM patients.

Our framework demonstrates the power of a multi-scale, segmentation-free self-supervised learning model. By training our model on cells we were able to characterize rare and novel cell subpopulations, and by training our model on LTMEs, we were able to capture complex tissue-wide spatial interactions. Our model is also scalable and reusable, and future studies conducted using larger datasets and additional markers could yield findings with greater predictive power. By rapidly generating hypotheses that can be validated with in vitro experiments, our framework could help researchers identify potential therapeutic targets.

## Methods

### Preprocessing of IMC image data

Our dataset included 389 IMC histopathology images from 185 patients, 139 of which had high-grade gliomas, and 46 had brain metastasis (BrM) tumors originating from various primary sites, including breast (11), lung (19), melanoma (7), ovarian (2), colorectal (1), gastric (1), gastro-intestinal (1), and bladder (2). We had survival data for 109 glioma patients and 45 BrM patients. The bulk of images of patient cores, although we also had margin samples from 41 of our BrM patients, and tumor-adjacent normal samples from 18 of our glioma patients. All core and margin slides were used for training and inference, but the tumor-adjacent normal slides were not used. The dataset also included a total of 1,163,362 cells that were segmented and classified using a supervised lineage assignment approach; for each cell, the cell type and boundary information was provided.

The images provided on Zenodo contained a total of 24 de-noised markers. The glioma and BrM images were stained for different markers and thus had different channels. In order to simultaneously train on both glioma and BrM data, we consolidated all channels into a multi-channel image. We later obtained an additional 13 raw marker channels from the lab, which had not been de-noised. To process these markers, we applied k-means clustering to determine a pixel intensity cutoff, removing pixels below the cutoff. We then used median filtering to further reduce noise by replacing each pixel with the median value of its neighboring pixels. We measured performance by applying our denoising algorithm to the CD94 channel and comparing the results with the CD94 channel denoised by the lab by calculating the Structural Similarity Index Measure (SSIM) score.

### Generating and normalizing tiles

To generate SCME tiles, we used the provided cell boundary data to generate 16-micron square tiles centered on individual cells. We chose 16-microns as the average tumor cell in our dataset is between 10-20 microns in diameter, and a 16-micron tile size strikes a balance between capturing most cells within the tile boundary and not capturing too much information from outside the cell boundary.

To analyze LTMEs, isolated relevant regions of the image by normalizing the image’s pixel value range to between 0 and 1, and selected regions where the average pixel value is above 0.2, and used those regions to generate our 64-micron square tiles. Because marker intensity across different channels can vary due to biological variations and imaging artifacts, we performed channel-wise normalization by sampling 10,000 random tiles from the dataset to calculate the mean and standard deviation for each channel. During training, input tiles were normalized using these statistics to ensure consistent scaling across the dataset.

Data augmentation was applied to reflect biological and technical variations. This consisted of random rotations (0 to 180 degrees), horizontal flips, random cropping (scaling from 50 to 100%), and channel intensity augmentation. For SCME tiles, our random cropping scaled from 80 to 100% to ensure that in most cases the entire cell remains inside the tile. To expedite training and ensure a diversity of cell types for our model training, we generated SCME tiles for 100 randomly sampled cells of each cell type from each image. After processing, this resulted in 230,953 SCME tiles for training, and 923,960 SCME tiles for inference. For LTMEs we generated 25,671 tiles, all of which were used for both training and testing.

### Model architecture and training

Our model consists of a Vision Transformer (ViT) encoder and a lightweight ViT decoder. The encoder is a ViT-Large with 24 transformer blocks, a hidden size of 1024, and 16 multi-head attention heads. The decoder has 8 blocks and a hidden size of 512. To account for spatial information, a 2D sine/cosine positional embedding is applied to the input patches. During model training, both our 16-micron SCME tiles and 64-micron LTME tiles are resized to 224×224 pixels and split into 256 patches of 14×14 pixels each. Each image patch is encoded by a convolutional neural network (CNN) with positional encodings are added to the patch embeddings. The model masks 75% of patches and uses them as reconstruction targets for the decoder, while the remaining patches are fed into the transformer encoder.

The encoded patches are then combined with masking tokens and restored to their original positions, followed by an additional positional embedding. These are then passed to a lightweight transformer decoder to predict the masked patches. Our SCME model was then trained on 230,953 cell tiles for 600 epochs, and our LTME model was trained on all 25,671 LTME tiles for 1600 epochs. The loss is computed using mean squared error (MSE) by comparing the predicted masked patches with the original patches **(Suppl. Fig. 1a)**, and our trained model is used to perform image reconstruction for our LTME and SCME tiles **(Suppl. Fig. 1b-c**).

### Clustering and visualization

To cluster tiles, we tested clusters between 3 and 80. Because the elbow plot did not yield a distinct elbow point, we performed clustering at 10, 20, 30, 40, 50, and 60 centroids. We settled on 40 centroids for both LTMEs and cells, as it enabled us to generate a diverse mix of clusters while minimizing the number of redundant clusters, which we assessed by plotting cell type composition and marker expression on a normalized bar chart for each cluster, and visually inspecting for similar clusters.

To validate our SCME clusters, we first created a heatmap comparing marker expression to the cell type annotations from the original dataset by calculating the mean pixel intensity for each cell tile, aggregating by cell type, then applying z-score normalization to standardize across cell types. We then repeated this with clusters, generating a heatmap showing the normalized expression of each marker by cluster **(Suppl. Fig. 2)**. We used a UMAP to our LTME and SCME tiles, in which each dot represents a tile’s feature embedding.

Lastly, we generated bar plots showing cell and marker composition for each cluster. The cell composition bar plot shows the percentage of different types of cells within a cluster. To create our marker composition barplot, we calculated marker expression by taking the mean pixel intensity for each channel in each tile and performing min-max scaling (for LTMEs, we removed outliers by only taking values within the 2nd and 98th percentile of marker values for each channel across samples) and then calculated the relative percentage of marker expression for each cluster.

### Survival analysis

Each patient has a different composition of SCME and LTME tiles. We refer to tile concentration (or TC) as the number of tiles a patient has for a specific LTME cluster, divided by their total number of tiles. For each cluster, we can then compute a median TC across patients. This lets us split patients into a high TC group if they exceed the median TC for a cluster, and a low TC group if they have equal or less than median TC for a cluster. We used Kaplan-Meier to plot survival curves for both groups, and calculated the p-value using a log rank test. To address rare clusters, we re-performed the tests with cutoffs at the 25th and 10th percentile for TC. To validate significant clusters further, we used bootstrapping with 1,000 resamples to calculate c-index (concordance index) scores, which measures the ability of a model to correctly rank the survival times of individuals, and does not require set cutoffs. A c-index score of 0.6 means that a model is able to correctly rank patient survival times for 60% of all possible patient pairs, and indicates moderate predictive power. Because our c-index model is ranking for increased survival, values under 0.5 suggest a relationship with decreased survival, so we subtract our c-index from 1 in cases where c-index is below 0.5 and the c-index coefficient is negative. For example, a score of 0.4 is equivalent to a score of 0.6.

### Data availability

The high dimensional IMC images used in this study, as well as the patient survival data, are available at this link: https://zenodo.org/records/7884599. Data used for scRNA-seq and snRNA-seq analysis is available from the Gene Expression Omnibus repository, accession codes <u>GSE131928</u> and <u>GSE174554</u>.

### Code availability [WIP]

The code used for training our model is available at these Github links: for our <u>LTME pipeline</u>, and our <u>SCME</u> <u>pipeline</u>. The code used to perform downstream analysis are available as Colab notebooks:

● LTME composition map and UMAP
● SCME composition map and UMAP
● LTME L2 and B cell survival analysis

## Supplementary

**Supplementary Fig. 1:**
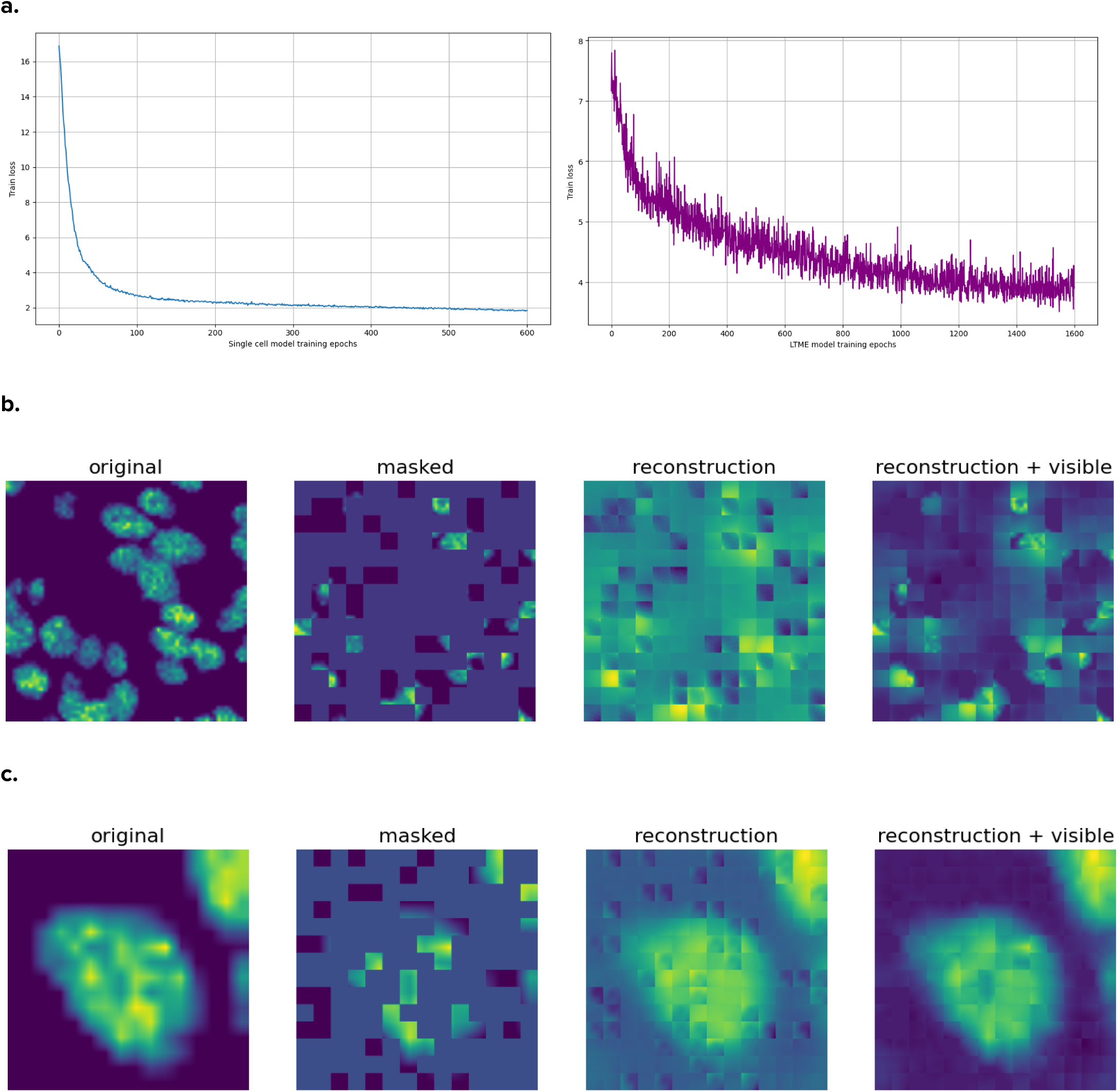
**a.** (Top) Mean squared error training loss for our SCME model and (bottom) MSE training loss for our LTME model. **b.** Image reconstruction performed on LTME tiles, using our trained model. **c.** Image reconstruction performed on SCME tiles.

**Supplementary Fig. 2:**
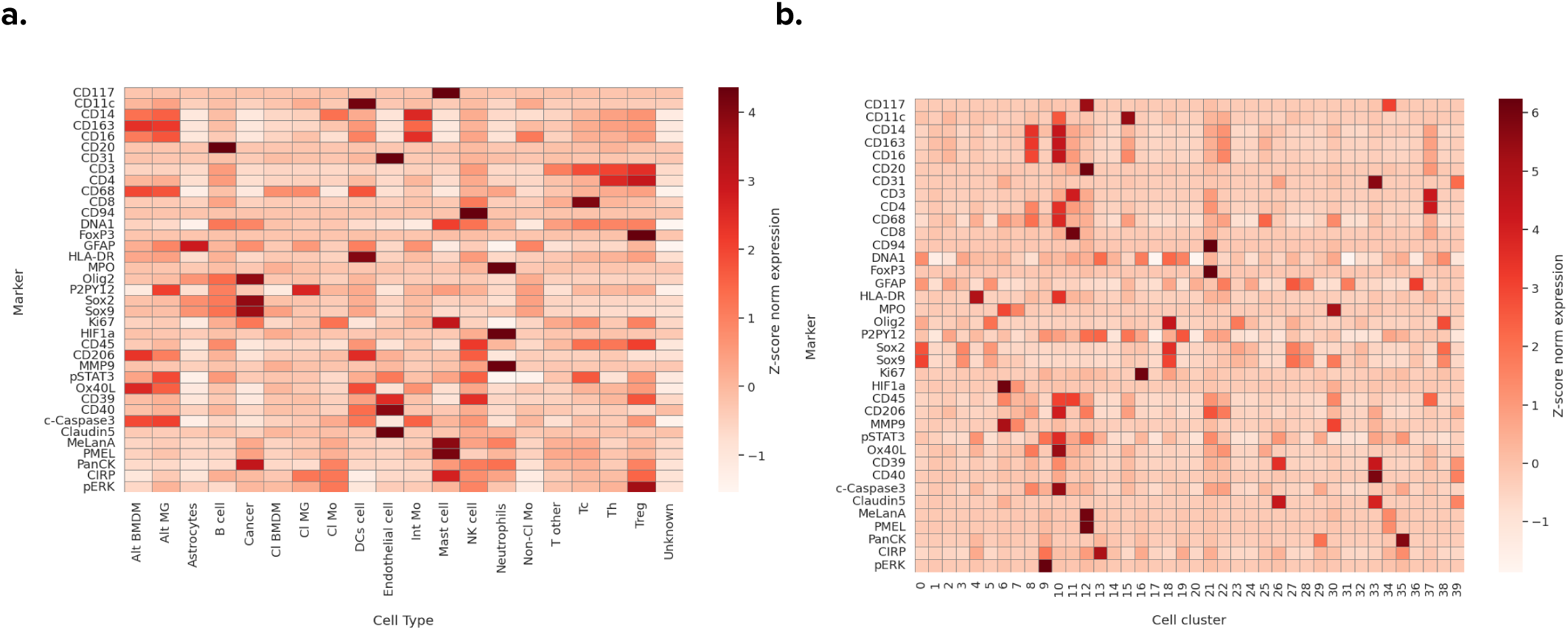
**a.** Heatmap comparing the labeled cells in the dataset, which were classified using a supervised lineage approach, to their summed and normalized marker expression. **b.** Heatmap comparing our cell clusters to their normalized marker expression.

**Supplementary Fig. 3:**
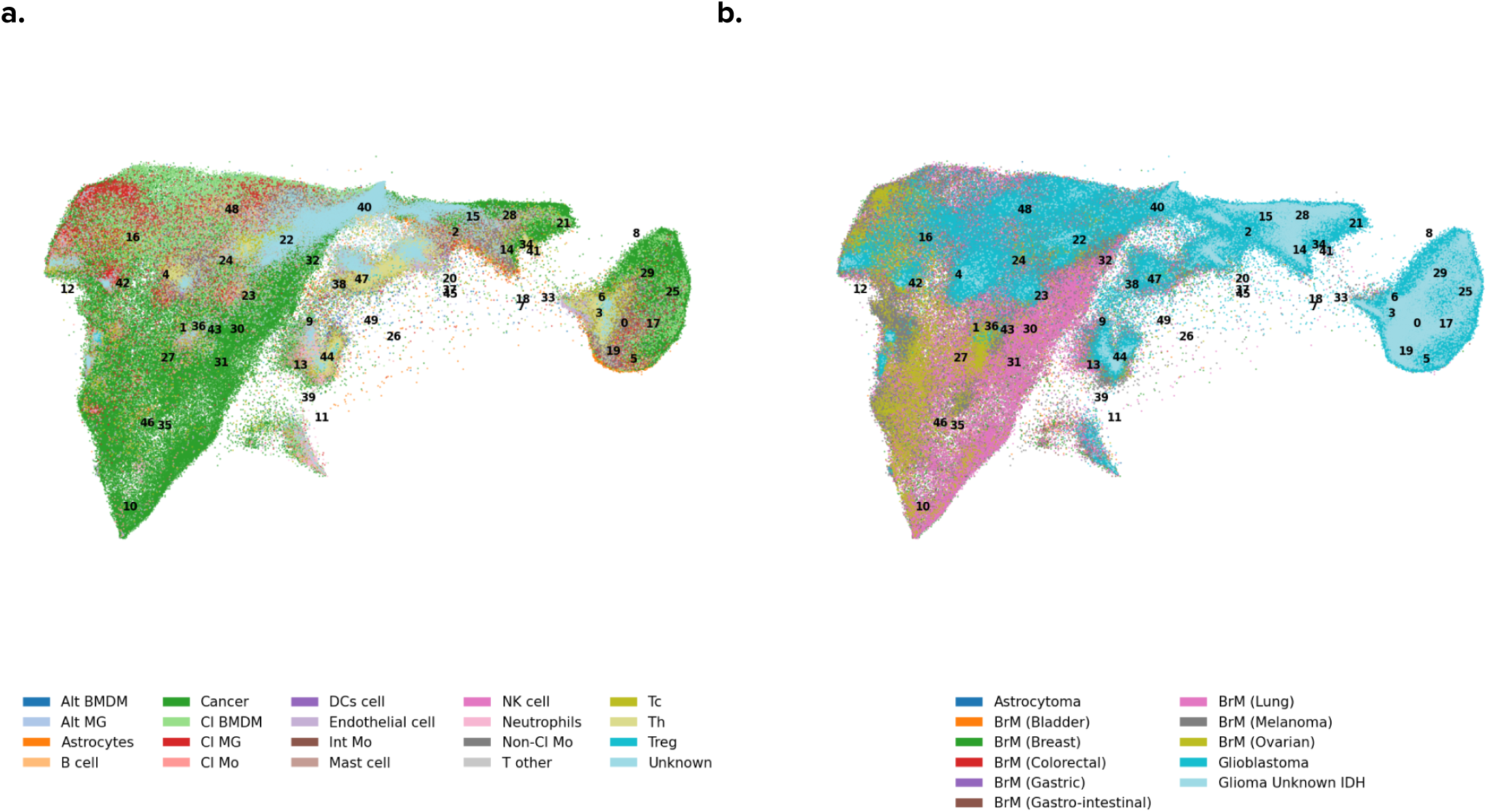
Self-supervised representation learning on SCME. **a.** UMAP, where each dot represents an SCME tiles’ feature embedding, colored by cell type to show that cell types tend to cluster together. **b.** UMAP is colored by tumor type, BrM cells are colored by the primary tumor’s origin.

**Supplementary Fig. 4:**
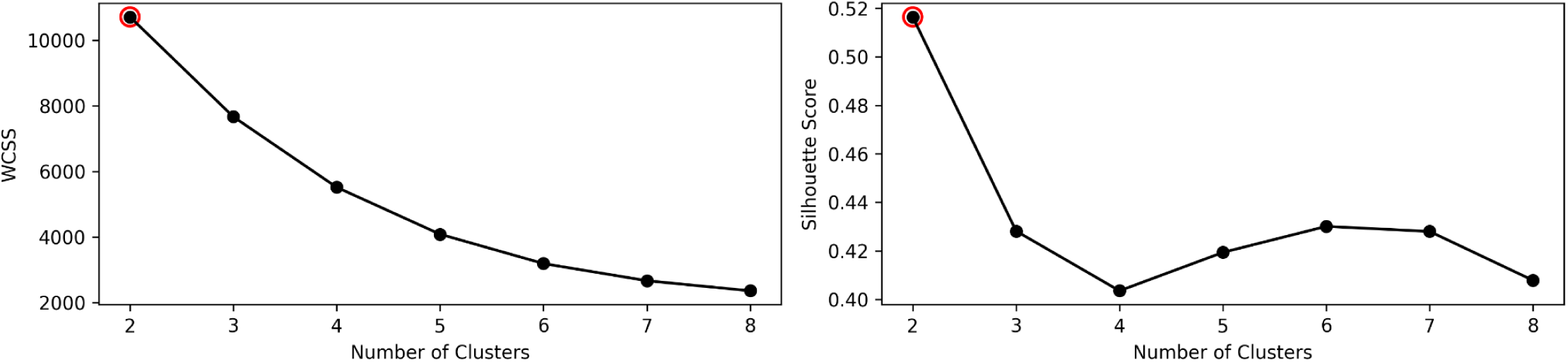
Silhouette and elbow plot to determine the optimal number of B cell clusters.

**Supplementary Fig. 5:**
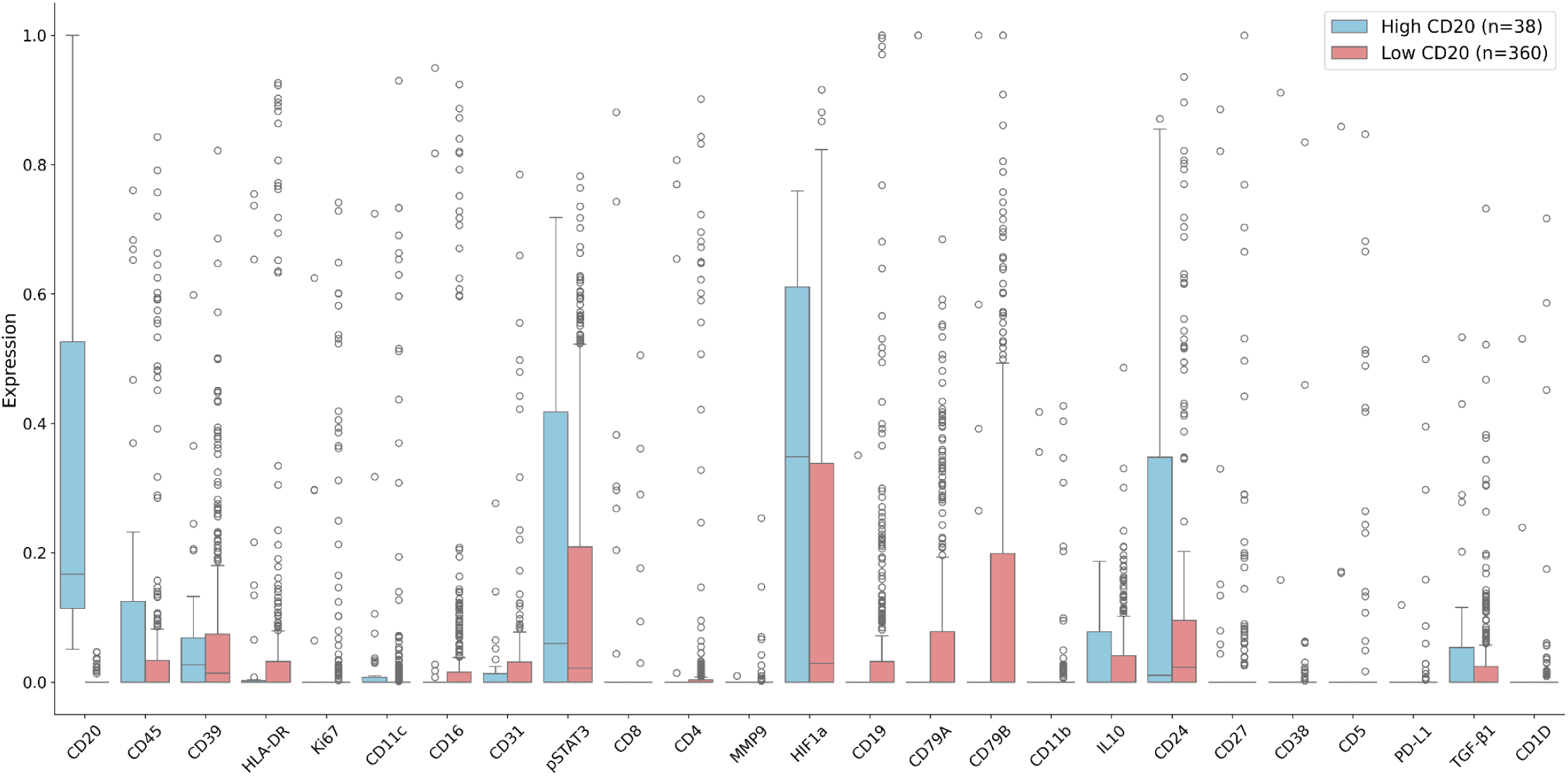
Boxplot showing the difference in marker expression between CD45+ and CD45- B-cells from our external dataset.

**Supplementary Fig. 6:**
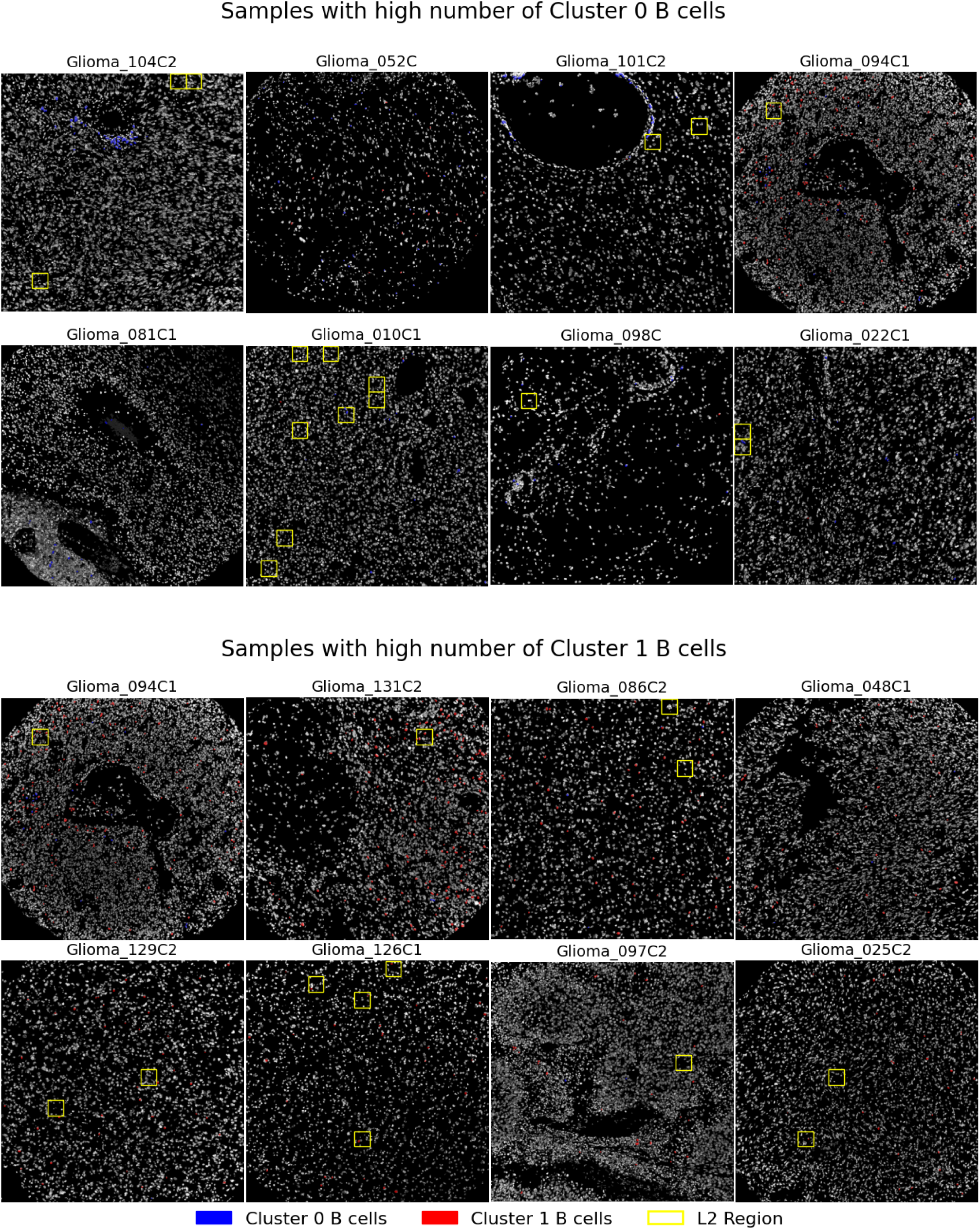
DNA1 channel colored by Cluster 0 and Cluster 1 B cells, showing that while C0 B cells are often found near perivascular regions, L2 (and Cluster 1 B cells) are typically found further away from perivascular regions.

**Supplementary Fig. 7:**
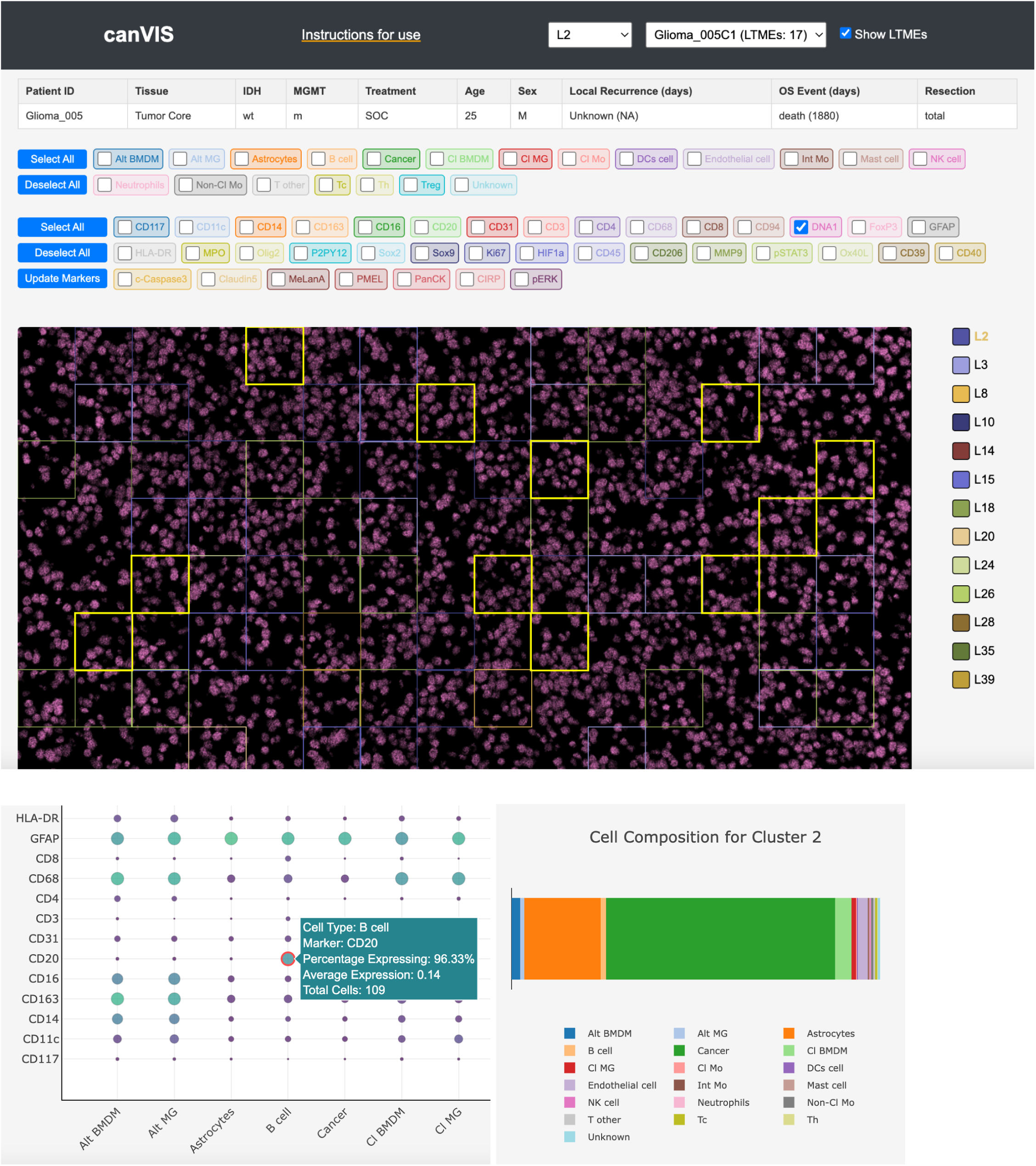
Our custom web app enables in-depth characterization of marker expression and spatial features within an LTME of interest and is included as part of this publication.

## Acknowledgements

We would like to thank Dr. Logan Walsh and team at the Quail-Walsh Lab for their support with our research, as well as for providing the additional 13 IMC markers used in our analysis. We would also like to thank Dr. Benjamin G. Neel at New York University Langone Health for providing helpful comments on our manuscript.

## Notes

https://zenodo.org/records/7884599

### Competing Interest Statement

The authors have declared no competing interest.

